# Source of genome-wide deleterious variation in a global cattle cohort

**DOI:** 10.64898/2026.09.07.749866

**Authors:** Junxin Gao, Martijn F. L. Derks, Job G. C. van Schipstal, Ying Liu, Etske Bijl, Catarina Ginja, Juha Kantanen, Nasser Ghanem, Donald R. Kugonza, Mahlako L. Makgahlela, Martien A. M. Groenen, Henk Bovenhuis, Richard P. M. A. Crooijmans

## Abstract

**Background:** Identifying deleterious DNA changes underpins efforts to improve animal health, welfare, and sustainable breeding. In cattle, current variant prioritization focuses on coding changes, uses single annotation types, and gives limited resolution in non-coding sequence.

**Results:** We developed BovCADD (bovine Combined Annotation-Dependent Depletion), a nucleotide-level deleteriousness score for substitutions in *Bos taurus* and *Bos indicus*, combining evolutionary constraint, sequence context, epigenetic and regulatory annotations, and gene and protein features. A logistic regression model trained on 41.9 million high-frequency derived alleles from about 3,700 cattle, contrasted with context-matched simulated variants, scored all 8.1 billion possible substitutions. BovCADD distinguished known pathogenic variants from background variation, discriminated among variants within the same consequence class, and scored intronic and intergenic sites. Aggregating scores identified genes carrying rare deleterious variation and revealed elevated genetic load at trait-relevant loci and in bottlenecked, intensively selected populations.

**Conclusions:** BovCADD provides the first genome-wide, nucleotide-resolution measure of deleteriousness in cattle, extending variant interpretation into non-coding sequence and linking variant-level prioritization with population-level patterns of mutational burden. Precomputed scores for all substitutions are publicly available.

## INTRODUCTION

Interpreting the effects of naturally occurring genetic variation in cattle is fundamental for improving animal health and welfare, accelerating genetic gain for dairy and beef production, and safeguarding diversity in both commercial and indigenous populations. Establishing variant–effect links in bovine remains challenging: most commercial and biological trait architectures are highly polygenic, long-range linkage disequilibrium can blur causal signals and confound fine-mapping [1, 2]. Functional evidence uneven across tissues, developmental stages, environments, and breeds, leaving large fractions of the bovine genome without clear interpretability, especially on regulatory and noncoding regions [3]. Routine variant prioritization in bovine genomics still often relies on single evidence streams, such as evolutionary conservation, predicted coding consequence, or association statistics from GWAS/QTL studies [4–8], each of which captures only a limited facet of biological impact and provides incomplete coverage of the noncoding genome.

Human genetics has demonstrated that combining heterogeneous annotations into a single, calibrated deleteriousness score can improve the discovery and interpretation of functional variation in the whole genome [9–11]. However, predictors trained in humans cannot be directly applied to cattle. The species differ in genome organization, regulatory landscapes, regions under selection, population structure and demographic history. Reflecting this, Combined Annotation Dependent Depletion (CADD)-style frameworks have been developed for multiple non-human species, including mouse (mCADD) [12, 13], pig (pCADD) [14], and chicken and turkey (chickenCADD and turkeyCADD) [15], and underscore the broader utility of species-specific scores for studying functional burden and genetic load in managed and wild animal populations [16].

The importance of domesticated cattle (*Bos taurus* and *Bos indicus*) in human nutrition, agricultural productivity and socio-economic systems underscores the need for species-specific deleteriousness scores [17–19]. Ongoing pressures from climate change, emerging infectious diseases, and resource constraints further increase the value of such tools. These tools can prioritize variants with plausible biological effects and quantify deleterious burden under inbreeding or reduced effective population size [16, 20]. A bovine deleteriousness score can therefore act as a practical “bridge” between association signals and underlying biological mechanisms: it can rank candidate variants within large GWAS/QTL intervals characterized by extensive linkage disequilibrium, including putative noncoding variants in promoters, enhancers, and other regulatory elements for functional follow-up [9, 14, 21]. When combined with cattle functional genomics resources, such as tissue-specific regulatory annotations, expression quantitative trait loci (eQTLs), and chromatin accessibility maps, it can help highlight variants most likely to perturb gene regulation in relevant tissues [22–24]. At the population level, CADD scores have been incorporated as prior information in genomic prediction [13, 25], detection of regions under selection [26], introgression and local ancestry analyses [27], and trait and genetic load estimation [12]. These applications help distinguish likely causal variants from linked hitchhiking variants. They also enable comparisons of deleterious variation across breeds, populations, and environments. In applied contexts, such scoring can accelerate the identification of candidate causal variants for Mendelian disorders [9, 28] and enable monitoring of deleterious alleles to reduce carrier-by-carrier mating risk and inform breeding strategies [29].

This study aimed to develop a bovine-specific framework to predict the deleterious effects of genetic variation across the genome. We introduce BovCADD, a bovine nucleotide-level deleteriousness score for *Bos taurus* and *Bos indicus* that integrates evolutionary constraint, local sequence context, bovine regulatory/epigenetic annotations, and gene/protein features into a single genome-wide metric. Following the CADD paradigm [9–11, 15], we trained a supervised machine learning model using 41.9 million high-frequency “proxy-benign” derived alleles contrasted with 41.9 million simulated single nucleotide variants (SNVs). We then precomputed scores for all 8.1 billion possible single-nucleotide substitutions on the widely used ARS-UCD1.2 assembly and newer bovine assemblies (ARS-UCD2.0).

BovCADD was validated through enrichment analyses in functional annotations and by benchmarking its performance against (i) known deleterious and functional variants from the Online Mendelian Inheritance in Animals database (OMIA) [28], (ii) an established cattle-specific functional framework, the Functional-And-Evolutionary Trait Heritability (FAETH) [5], and (iii) the distribution of BovCADD scores integrated across multiple annotation types [30]. Beyond validation, we further explore several applications of BovCADD, illustrating its utility by profiling genome-wide distributions of rare deleterious variants, trait-relevant variant burden using the Animal Quantitative Trait Loci Database (Animal QTLdb), and putative genetic load across global cattle populations [31–34]. Together, these results establish BovCADD as a broadly applicable resource for prioritizing both coding and noncoding variation in cattle. We release genome-wide score tables for multiple reference assemblies, along with the trained model, feature definitions, and reproducible code, to support transparent reuse in fine-mapping, functional follow-up, and breeding applications.

## RESULTS

### Implementation of BovCADD

We contrasted variants likely tolerated in cattle populations with sequence context–matched simulated mutations, following the CADD framework adapted to the bovine genome (Fig. 1A,1B) [9, 35]. Derived alleles were inferred by comparing *Bos taurus* and a closely related outgroup (*Ovis aries*) genomes to an ancestral sequence reconstructed from the 43-eutherian-mammal Ensembl EPO multiple alignment (v113). Sites at which the 1000 Bull Genomes Project [36] and OPTIBOV datasets [37] exhibited a derived allele frequency (DAF) ≥ 95% were designated as “proxy-benign”, yielding 41.9 million variants that are fixed or nearly fixed on the bovine lineage. To simulate an equivalent number of de novo mutations, we applied an empirical sequence-evolution model with CpG-specific substitution rates and mutation rates locally estimated on 1 Mb scales, following the human CADD procedure [9].

**Fig. 1.**
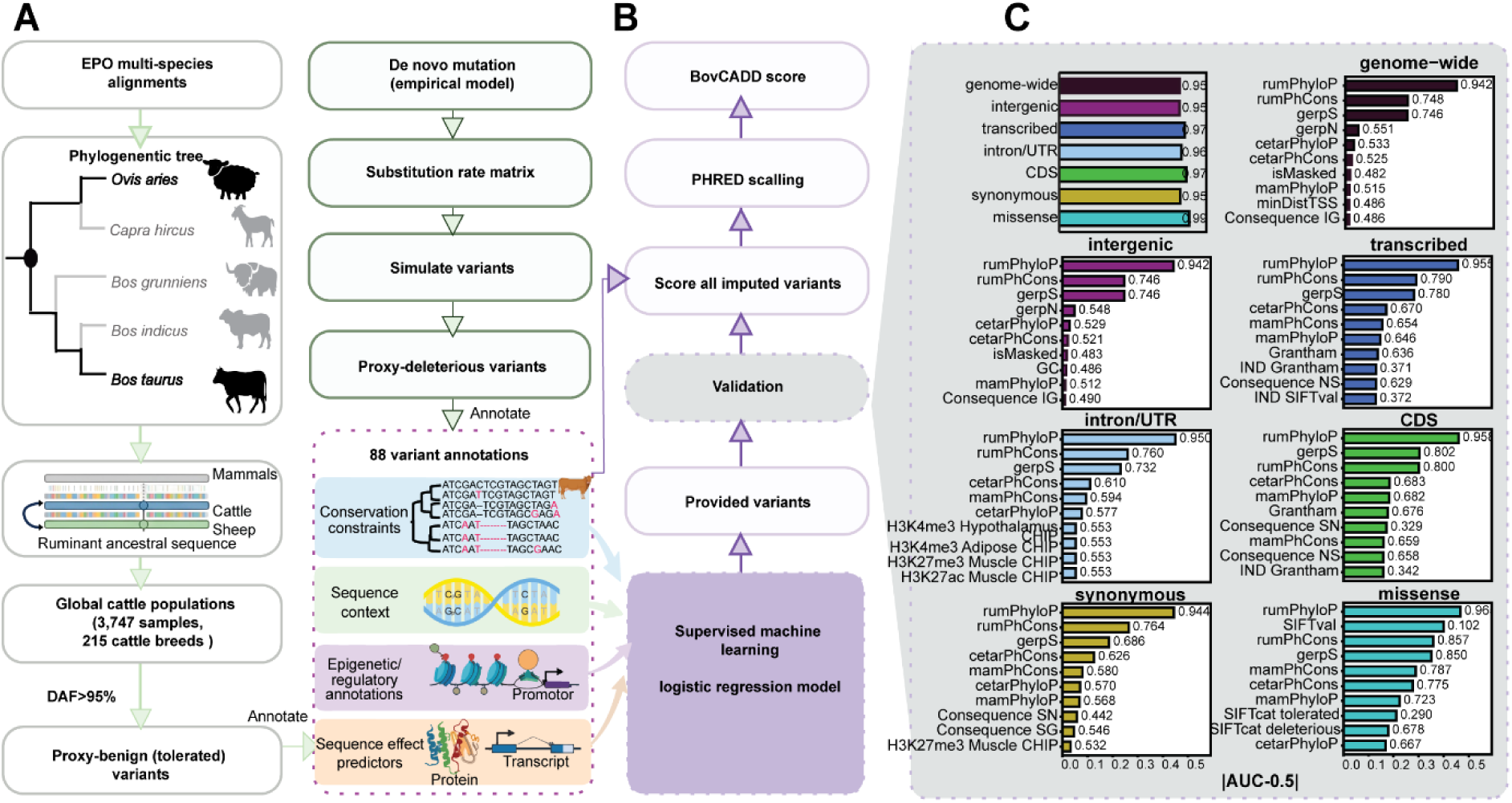
The BovCADD framework. **(A)** Training data. Ancestral sequence was reconstructed from the Ensembl EPO multi-species alignment, with *Ovis aries* as outgroup. Derived alleles at DAF > 95% across 3,747 cattle from 215 breeds were designated proxy-benign; an equal number of proxy-deleterious variants was simulated under an empirical de novo mutation model with CpG-aware substitution rates. Both sets were annotated with the same 88 features, spanning conservation, sequence context, regulatory annotations, and sequence-effect predictors. **(B)** Scoring and validation. An L2-regularized logistic regression model was trained to separate the two classes. All possible single-nucleotide substitutions in ARS-UCD1.2 were scored and converted to PHRED-scaled BovCADD values by rank. Performance was assessed by five-fold cross-validation and by external validation on independent variants from Online Mendelian Inheritance in Animals (OMIA; https://www.omia.org). **(C)** Single-annotation discrimination. For each annotation, the area under the receiver operating characteristic curve (ROC–AUC) was computed individually and is shown as |AUC − 0.5|, where 0 indicates no separation. The ten most discriminative annotations are given for seven variant classes: genome-wide, intergenic, transcribed, intron/UTR, coding, synonymous, and missense. The upper-left panel shows overall model ROC–AUC per class. Annotations are substantially inter-correlated, so these values are not additive.

For each candidate variant, we obtained functional annotations using Ensembl Variant Effect Predictor (VEP) [38] (ARS-UCD1.2), UCSC genome (bosTau9) [39], and bovine epigenetic and regulatory datasets (ATAC-seq and ChIP-seq peaks, transcription-factor (TF) motif metrics) into a uniform schema [40–42]. The feature set comprised: (i) evolutionary constraint, including Genomic Evolutionary Rate Profiling (GERP) scores [43] and multi-scale conservation metrics (phastCons [44] and phylogenetic *P*-values score: phyloP) [6] derived from ruminant, cetartiodactyla, and mammalian alignments; (ii) regulatory context, defined by overlaps with ATAC-seq and ChIP-seq peaks (CTCF and histone marks H3K27ac, H3K27me3, H3K4me1, H3K4me3) across adipose, cerebellum, cortex, hypothalamus, liver, lung, muscle, spleen, rumen tissues, and commonly profiled cell lines (Table S1) [41], together with transcription factor (TF)-motif distribution scores [45], chromatin states, and distances to transcription start sites (TSS); (iii) local sequence context, including reference/alternate nucleotides, transition/transversion class, GC and CpG content, DNA-shape features (HelT, MGW, ProT, Roll), and proximity to transcriptional boundaries; and (iv) gene/protein annotations, including VEP consequence categories [38], protein-domain and amino-acid substitution properties (original/new amino acid, Grantham score [46], Sorting Intolerant From Tolerant (SIFT) [47], and relative positional features within cDNA, CDS, and protein coordinates. The resulting variant-by-annotation matrix comprised 88 features (Table S2) across 83.8 million labelled variants in balanced observed/simulated subsets.

As observed in previous CADD models, conservation-based annotations consistently ranked among the top predictors across all genomic regions examined (genome-wide, non-cDNA, cDNA, non-CDS, CDS, synonymous, and missense), with predictive importance varying across these contexts [14, 15] (Fig. 1C, Data S1). Ruminant lineage-specific conservation scores (rumPhyloP, rumPhCons) consistently outperformed their broader Cetartiodactyla- and mammal-wide counterparts (cetarPhyloP/cetarPhCons and mamPhyloP/mamPhCons, respectively). In contrast, chromatin- and tissue-specific annotations (e.g., H3K27ac_Muscle, H3K4me3_Hypothalamus, TFmotif) showed receiver operating characteristic area under the curve (ROC–AUC) values close to the null expectation of 0.5 in most regions, with modest increases restricted to non-coding contexts (non-CDS, synonymous). SIFT-based annotations displayed region-specific behavior, showing negligible discriminative power outside coding sequence but the strongest single-feature signal within missense variants (SIFTval AUC = 0.102; |AUC − 0.5| = 0.398). Accordingly, the best-performing individual annotations for missense variants were conservation- and deleteriousness-based metrics such as PhyloP and SIFT; missense variants comprised 0.71% of the training set, of which 99.96% and 97.35% had defined rumPhyloP and SIFT values, respectively. Across five-fold cross-validation, BovCADD achieved an AUC of 0.955 (s.d. 6.5 × 10⁻⁵) and accuracy of 0.911, with near-identical results at both regularization strengths tested (C = 0.1 and C = 10; Table S3). Correlation analyses revealed substantial inter-dependence among annotations; because many features captured overlapping information, their predictive contributions were partly redundant rather than additive (Data S2). Collectively, these analyses demonstrate substantial biological differentiation between observed and simulated variants across the 88 annotations and confirm that a linear model captures most of this information.

We then trained a supervised machine-learning model (an L2-regularized logistic regression [48]) from the 88 annotations, supplemented with a limited set of interaction terms (Data S3). Predictions were averaged across cross-validation models and used to score all 8.1 billion possible single-nucleotide substitutions in the bovine reference genome (ARS-UCD1.2). Raw scores were transformed into Phred-scaled BovCADD values based on their rank among all possible substitutions, ranging from 1 to 99 [49]. Variants in the top 10%, 1%, 0.1% and 0.01% of scores corresponded to BovCADD thresholds ≥ 10, ≥ 20, ≥ 30, and ≥ 40, respectively.

### Genome-wide properties of BovCADD

The distribution of BovCADD scores across all possible single-nucleotide substitutions was quantified and their functional consequences were summarized (Fig. 2A). BovCADD scores were the highest for predicted loss-of-function variants, with stop-gained substitutions showing the greatest mean score (30.8), followed by canonical splice-site (22.3) and missense variants (16.5). In contrast, intergenic substitutions had the lowest average scores (3.4). Despite this enrichment, a substantial fraction of high-scoring variants was noncoding: 44.95% of all substitutions with BovCADD ≥ 20 fell outside missense, stop-gained, stop-lost, or splice-site categories. Conversely, 65.59% of potential missense variants and 8.94% of potential stop-gained variants had BovCADD scores < 20.

**Fig. 2.**
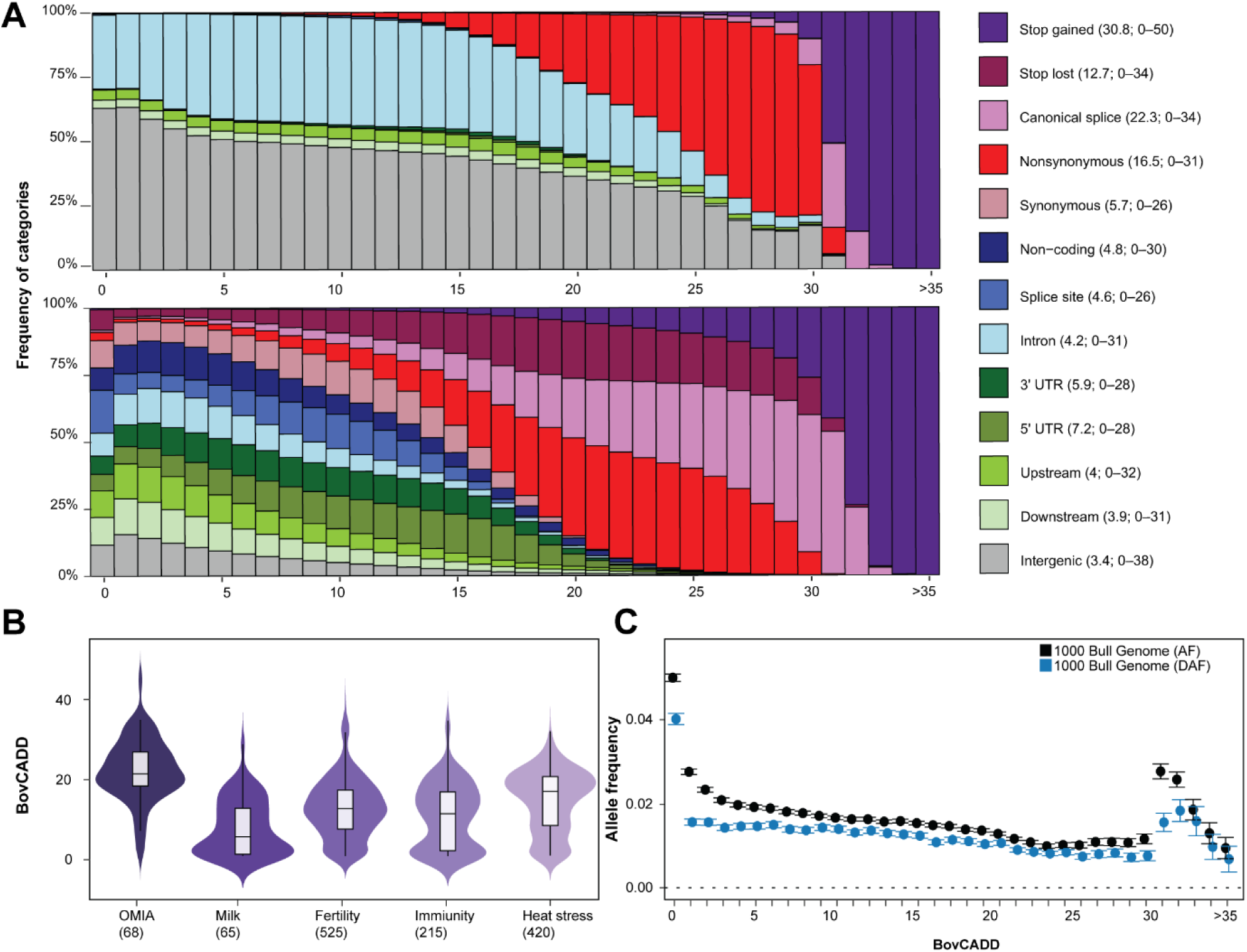
Relationship between BovCADD scores and genome-wide variant properties. **(A)** Association between BovCADD scores and categorical variant consequences. Top, proportion of substitutions annotated with each functional consequence within BovCADD score bins. Bottom, the same proportions normalized by the total number of variants in each consequence category. Consequences were annotated with the Ensembl Variant Effect Predictor (VEP). Scores above 35 collapsed into a single bin (> 35) owing to low variant counts; all variants in this bin were stop-gain. **(B)** The OMIA set comprises all 68 catalogued variants irrespective of consequence (https://www.omia.org); the trait-associated sets are restricted to protein-altering variants, defined as stop-gain, missense, start-lost, and splice-site consequences. Milk-production variants were obtained from the 1000 Bull Genomes Project (Run 9) and OPTIBOV. Variants associated with fertility (*EIF4EBP3*, *ATP10A*), immunity (*IGLL1*, *TLR4*), and heat-stress–related traits (*HSPA12B*, *HSF1*) were identified from Ensembl gene annotations (July 2023 release; https://jul2023.archive.ensembl.org), with functional consequences annotated using Ensembl VEP. **(C)** Relationship between BovCADD scores and allele frequency. Mean derived allele frequency (DAF) and mean allele frequency (AF) are shown across BovCADD score bins for variants from the 1000 Bull Genomes Project (Run 9). Error bars are 95% confidence intervals (±1.96 standard error).

Within functional classes, BovCADD further separated variants by score (Fig. 2B). OMIA catalogues variants with established causal roles in Mendelian disorders and inherited traits in cattle [28], providing a set of high-confidence functional variants independent of our training data. These scored highly overall (mean = 22.37; Table S4), exceeding protein-altering variants associated with specific production and health traits, which spanned a wide range and differed by trait: heat tolerance (14.51), fertility (12.52), immunity (10.53), and milk production (7.33) [50]. Such within-class resolution was absent or attenuated in commonly used alternatives, either through missingness (predictors restricted to protein-coding variants or to particular trait categories [47]) or through limited functional resolution, as with conservation scores [43], which cannot distinguish nonsense from missense substitutions at the same position. BovCADD therefore captures information both across and within functional categories, supporting genome-wide prioritization of potentially deleterious variants. Scaled BovCADD scores were next compared with patterns of genetic diversity. Scores correlated negatively with both DAF and allele frequency (AF) across the full spectrum in variants catalogued by the 1000 Bull Genomes Project [36] (Fig. 2C): high-scoring variants were progressively depleted among common alleles, consistent with purifying selection acting against deleterious mutations.

### BovCADD against known functional variants and trait-associated loci

We next evaluated the ability of BovCADD to prioritize functional and Mendelian disease–associated variants curated in OMIA. Across all bovine genomic sites, BovCADD discriminated pathogenic from benign variants with an AUC of 0.998 (Fig. 3A, Table S5). This performance exceeded that of individual annotation metrics, including phyloP (0.972), phastCons (0.884), GERP (0.892), Grantham (0.820) and SIFT (0.813), reflecting the added predictive value of integrating multiple annotation layers. Nearly all known variants underlying monogenic disorders and major production traits in cattle fell within the top 1% of BovCADD scores (range 1.43–45.78; mean 22.32) (Fig. 2B, Table S4). To characterize the genome-wide BovCADD score landscape, we contrasted scores between coding (mean = 9.51) and noncoding regions (mean = 3.75) (Fig. 3B). The highest-scoring OMIA variants were enriched for high-impact consequences: a stop-lost variant in *SDE2* (stop-lost mutation; associated trait: abortion due to haplotype HH6), *TMEM95* (stop-gain mutation; associated trait: male subfertility), *GATA6* (stop-gain mutation; associated trait: persistent truncus arteriosus), and *CEP250* (stop-gain mutation; associated trait: caprine-like generalized hypoplasia syndrome). Additional high-scoring variants included nonsense mutations in *CWC15* causing abortion due to haplotype JH1, missense mutations in *CYP26C1* associated with mandibulofacial dysostosis, pathogenic alleles in *MAP2K2* linked to skeletal–cardio–enteric dysplasia, missense variants in *ITGA3* responsible for de novo defects in an artificial-insemination sire, and splicing variants in *CCDC189* associated with asthenospermia. Regulatory variants in *ACAN* associated with Bulldog calf syndrome showed moderate BovCADD scores (6.62).

**Fig. 3.**
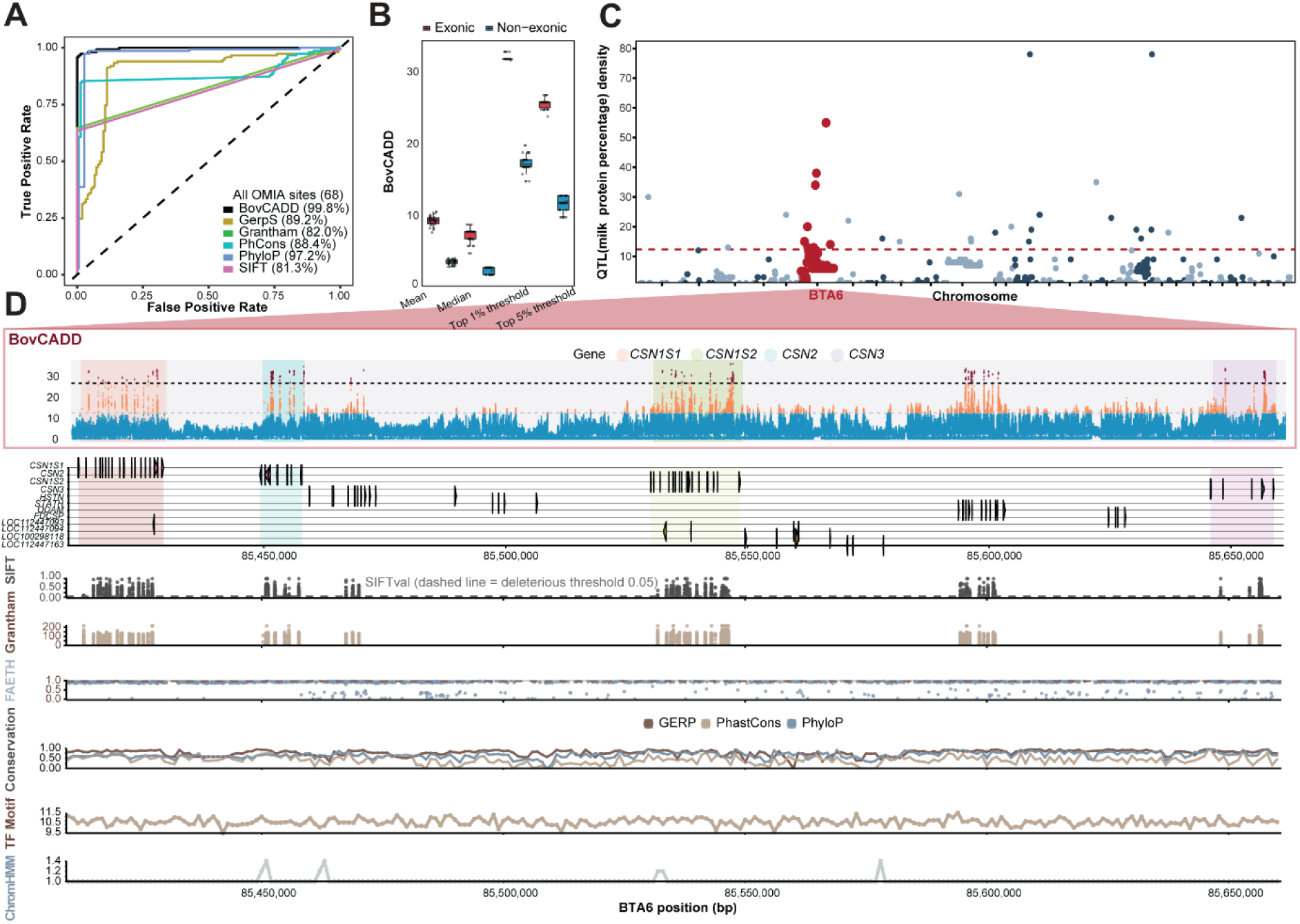
BovCADD scores prioritize functional genetic variation. **(A)** Performance of variant effect predictors in distinguishing pathogenic from benign variants. AUC compares BovCADD with genome-wide conservation- and impact-based metrics, including Genomic Evolutionary Rate Profiling score (GERP), PhastCons score, phylogenetic *P*-values score (phyloP), Sorting Intolerant From Tolerant score (SIFT), and Grantham score, using selected functional mutations from the OMIA database (*n* = 68). **(B)** Genome-wide distribution of BovCADD scores in exonic and noncoding regions. Points denote per-chromosome distributions; horizontal lines mark the median and error bars ±1.96 standard error. (**C)** QTL density for milk protein percentage based on the Animal QTLdb (1,213 QTL regions). The highlighted interval spans 85.42–85.46 Mb on chromosome 6, corresponding to a QTL region previously reported in study [51]. The dashed line indicates the top 5% QTL density threshold. **(D)** Representative locus showing BovCADD score distributions across a gene cluster associated with milk traits (85.42–85.46 Mb). Tracks show annotated protein-coding and long non-coding genes together with a subset of the 88 functional annotations, including SIFT, Grantham, conservation metrics, FAETH scores, transcription factor binding motifs, and ChromHMM chromatin states. Dashed lines mark the median top-5% BovCADD threshold across chromosomes, calculated separately for exonic (26) and non-exonic (12) sites.

To evaluate the genome-wide utility of BovCADD for functional variant prioritization within trait-associated regions, we used bovine milk protein percentage as a representative example. This trait is associated with 1,213 QTL regions in the Animal QTLdb (Fig. 3C) [30]. We then examined a well-defined trait-relevant gene cluster on chromosome 6 (85.54–85.62 Mb) [51], encompassing the core milk protein genes *CSN1S1*, *CSN1S2*, *CSN2*, and *CSN3*, as well as neighboring protein-coding and long noncoding genes within the broader QTL interval (Fig. 3D). Within coding sequences, BovCADD accurately prioritized amino acid-altering variants with strong deleterious signatures defined by SIFT and Grantham annotations (Fig. 3D). Within coding sequence, BovCADD prioritized amino acid– altering variants carrying deleterious signatures from SIFT and Grantham; in noncoding regulatory sequence, it combined conservation metrics (phyloP, GERP) with regulatory annotations to separate putatively functional variants from background variation (Fig. 3D). BovCADD therefore consolidates complementary signals from 88 annotations, supporting uniform prioritization across coding and noncoding sequence within trait-associated regions.

Finally, we compared BovCADD with the cattle-specific FAETH score [52] (fig. S1A, Data S4). Correlation was confined to the highest-ranking variants: within the top 10% of FAETH scores, median *r* = 0.47 across the autosomes, while all lower deciles fell between −0.07 and +0.08 (fig. S1B). Collectively, these results demonstrate that BovCADD is a powerful integrative framework for genome-wide functional variant prioritization. It complements existing cattle-specific prediction models, provides robust variant interpretation across coding and noncoding QTL domains, and supports downstream functional genetic studies and precision genomic breeding applications in cattle.

### Rare deleterious variation and gene-based burden

To quantify how deleterious variation accumulates across genes, we performed a genome-wide gene-based burden analysis using BovCADD on whole-genome sequence data from the 1000 Bull Genomes Project [36], following principles established in large-scale human studies of rare germline variation [53]. Because purifying selection preferentially removes damaging alleles, deleterious variants are expected to persist at low population frequencies (Fig. 2C); we therefore aggregated rare deleterious variants per gene and ranked genes by cumulative BovCADD burden. Four increasing thresholds (>20, >25, >30, and >35 rare deleterious variants per gene) identified 16,117, 10,064, 4,522, and 658 high-burden genes, respectively (Data S5). These genes were unevenly distributed across the genome (Fig. 4A), tracking variation in gene density across cattle chromosomes. As expected, genotype composition shifted progressively with threshold (Fig. 4B). Reference homozygotes predominated at every cutoff, while both alternative heterozygotes and alternative homozygotes declined as the threshold rose. Alternative homozygotes were the least represented class throughout, indicating that rare deleterious variants are maintained largely in the heterozygous state, masked from selection, and that homozygosity is selected against. Functional enrichment of high-burden genes recovered biologically coherent pathways, including cardiomyopathy- and disease-associated gene sets, cancer-related signalling cascades, cell–cell and cell–matrix adhesion, junction assembly, autophagy, and cell migration (Fig. 4C). This suggests that BovCADD burden mapping captures rare deleterious variation within functionally constrained networks.

**Fig. 4.**
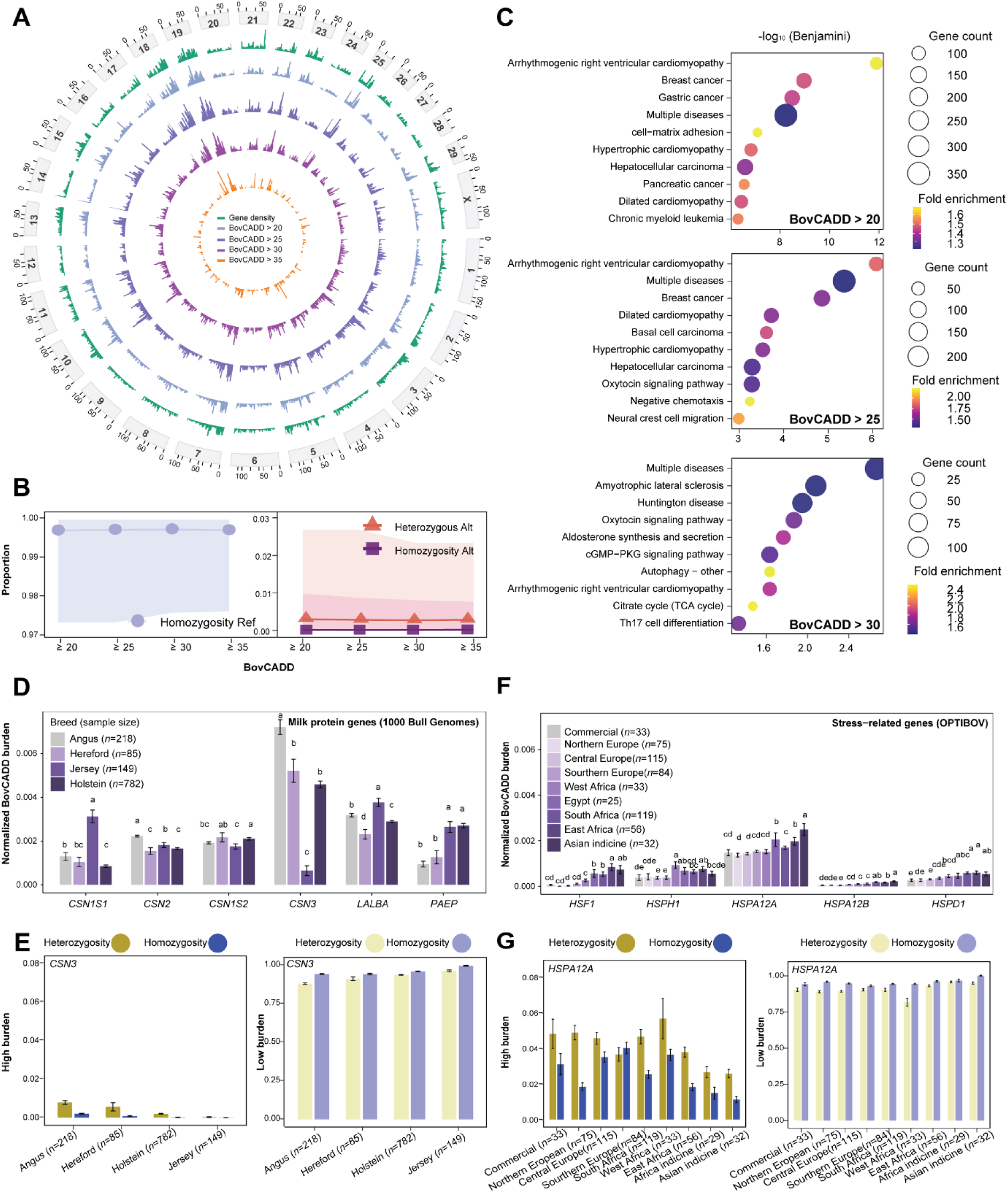
Genome-wide distribution and population-specific patterns of BovCADD gene-based burden. **(A)** Chromosomal distribution of high-burden genes. The circos plot shows genes exceeding four increasingly stringent BovCADD burden thresholds (>20, >25, >30, and >35 rare deleterious variants per gene). Gene density is shown as genes per Mb. **(B)** Genotype composition across burden thresholds. Proportions of reference homozygous (blue), alternative heterozygous (red), and alternative homozygous (purple) genotypes at each threshold. Each point is the mean proportion across all high-burden genes at that threshold; shaded regions are 95% confidence intervals. **(C)** Gene Ontology (GO) and Kyoto Encyclopedia of Genes and Genomes (KEGG) pathway enrichment analyses for genes exceeding BovCADD >20, >25, and >30 thresholds. Dot size indicates the number of genes per pathway, and color represents fold enrichment. Only pathways significant at Benjamini-adjusted *P* < 0.05 are shown. **(D)** Normalized BovCADD gene burden at major milk protein genes (*CSN1S1, CSN2, CSN1S2, CSN3, LALBA,* and *PAEP*) across four intensively selected commercial breeds (Angus, Hereford, Jersey, and Holstein–Friesian). Error bars indicate ±1.96 standard error. **(E)** Proportion of high- and low-burden variants at *CSN3*, stratified by heterozygous and homozygous state. Error bars indicate ±1.96 standard error. **(F)** Normalized BovCADD burden at stress-response genes (*HSF1, HSPH1, HSPA12A, HSPA12B*, and *HSPD1*) across global cattle populations. Error bars indicate ±1.96 standard error. **(G)** Proportion of high- and low-burden variants in stress-response genes across population groups, stratified by heterozygous and homozygous state. Error bars indicate ±1.96 standard error. Population definitions (**F,G**). Commercial breeds include Angus (6), Hereford (11), Jersey (6), and Holstein–Friesian (10). Northern Europe includes Western Finncattle (25), Northern Finncattle (25), and Eastern Finncattle (25). Central Europe includes Meuse Rhine Yssel (23), Groningen White Headed (21), Deep Red (24), Dutch Belted (23), and Dutch Friesian (24). Southern Europe includes Barrosã (28), Mertolenga (29), and Mirandesa (27). West Africa includes Muturu (23) and N’Dama (10). Southern Africa includes Afrikaner (40), Bonsmara (20), Nguni (40), and Ankole (19). East Africa and African indicine include Nganda (27), Karamojong (11), Kenana (8), Ogaden (7), and Goffa (3). Asian indicine includes Nelore (6), Brahman (14), Gir (5), and Boran (7).

To assess whether BovCADD gene-based burden captures deleterious loci with direct or indirect relevance to commercial and biologically important traits, two exemplar QTL-linked gene sets were analyzed: (i) six major milk-protein genes (*CSN1S1, CSN1S2, CSN2, CSN3, LALBA* and *PAEP*)[54]; and (ii) stress-response genes including members of the HSP70 family (*HSPA12A, HSPA12B), HSP60 (HSPD1), HSP110 (HSPH1)*, and the heat-shock transcription factor (*HSF1*) [33, 55–58]. For the milk-protein loci, we focused on four intensively selected commercial breeds (Fig. 4D,4E). Consistent with long-term directional selection and strong functional constraint on milk-protein genes, Holstein– Friesian cattle exhibited lower deleterious burden at *CSN1S1*, *CSN2* and *LALBA*, whereas Jersey cattle showed lower burden at *CSN1S2* and *CSN3*[54, 59]. *LALBA* encodes α-lactalbumin, a key regulatory component of lactose synthase that influences lactose synthesis and milk volume, while *CSN1S2* (αs2-casein) and *CSN3* (κ-casein) play critical roles in casein micelle structure and milk coagulation [60, 61]. Stratifying individuals by burden level and genotype state resolved this further: at *CSN3*, the locus showing the largest between-breed difference (Fig. 4D), Jersey cattle had the lowest proportion of high-burden variants and the highest proportion of low-burden variants in both homozygous and heterozygous classes (Fig. 4E), indicating that both recessive and additive deleterious variation has been reduced at this locus. In contrast, both Jersey and Holstein–Friesian cattle displayed relatively elevated burden at *PAEP* (*β*-lactoglobulin), a highly polymorphic whey protein gene previously shown to be subject to balancing selection in dairy populations, consistent with the maintenance of multiple functional alleles [62].

Burden at stress-response genes differed markedly among worldwide populations spanning diverse ecosystems, drawn from the 1000 Bull Genomes Project [36] and OPTIBOV datasets [37] (Fig. 4F,4G). European taurine breeds showed constrained burden distributions, whereas African indigenous and indicine breeds showed broader distributions and higher overall burden at several loci. Many African cattle carry admixed taurine–indicine ancestry [63], which contributes to this broader distribution. Stratifying individuals by burden level and genotype state, African and Asian indicine cattle carried the lowest proportions of high-burden variants and the highest proportions of low-burden variants in both homozygous and heterozygous states. This pattern is consistent with long-term adaptation to thermal and environmental stress, in which purifying selection has reduced deleterious variation in stress-response pathways [64, 65]. Together, these analyses show that BovCADD-based burden mapping characterizes genome-wide accumulation of deleterious variation and provides a framework for interpreting deleterious burden across complex traits.

### Genetic loads across global cattle populations

Genetic load reflects the cumulative burden of deleterious variation carried by an individual or population, and captures information distinct from association mapping, selection scans, or diversity-based analyses [16]. Whereas previous cattle genomic studies have focused largely on trait-associated loci and demographic history, BovCADD makes it possible to measure the retention of deleterious mutations genome-wide, and the efficacy with which purifying selection removes them, across diverse functional annotations [12, 16].

Cross-population comparisons of genetic load in cattle are complicated by population structure, genomic divergence between taurine and indicine subspecies, and widespread intersubspecific admixture [66–68]. Indicine and crossbred cattle carry elevated genome-wide sequence diversity, which inflates raw variant counts without a corresponding increase in functional deleterious burden. To enable standardized comparison, we stratified variants into low-burden (BovCADD < 5) and high-burden (BovCADD > 20) categories and quantified their proportions in homozygous and heterozygous states. Expressing genetic load as a proportion of each individual’s own variant set rather than as a count control for differences in baseline genomic diversity between populations.

Global patterns of BovCADD-based genetic load are summarized in Fig. 5A. We first examined the 32 taurine populations from the 1000 Bull Genomes Project represented by more than 20 individuals (*n* = 3,894; Table S6). Commercial breeds offer an informative system for examining deleterious variant accumulation, since sustained artificial selection for production traits and heavy use of elite sires can enrich deleterious variation through genetic hitchhiking and elevated relatedness [69]. Consistent with this expectation, dairy breeds carried the highest proportions of high-burden variants among commercial cattle in both homozygous and heterozygous states (Jersey 0.455% and 0.583%; Holstein 0.452% and 0.585%), with intermediate beef breeds (Angus 0.454% and 0.561%) and dual-purpose breeds lowest (Fig. 5B, fig. S2A, S2B). Dual-purpose breeds correspondingly showed the highest proportions of low-burden variants (Fig. 5C, fig. S2C, S2D), consistent with broader genetic diversity and less accumulation of deleterious alleles.

**Fig. 5.**
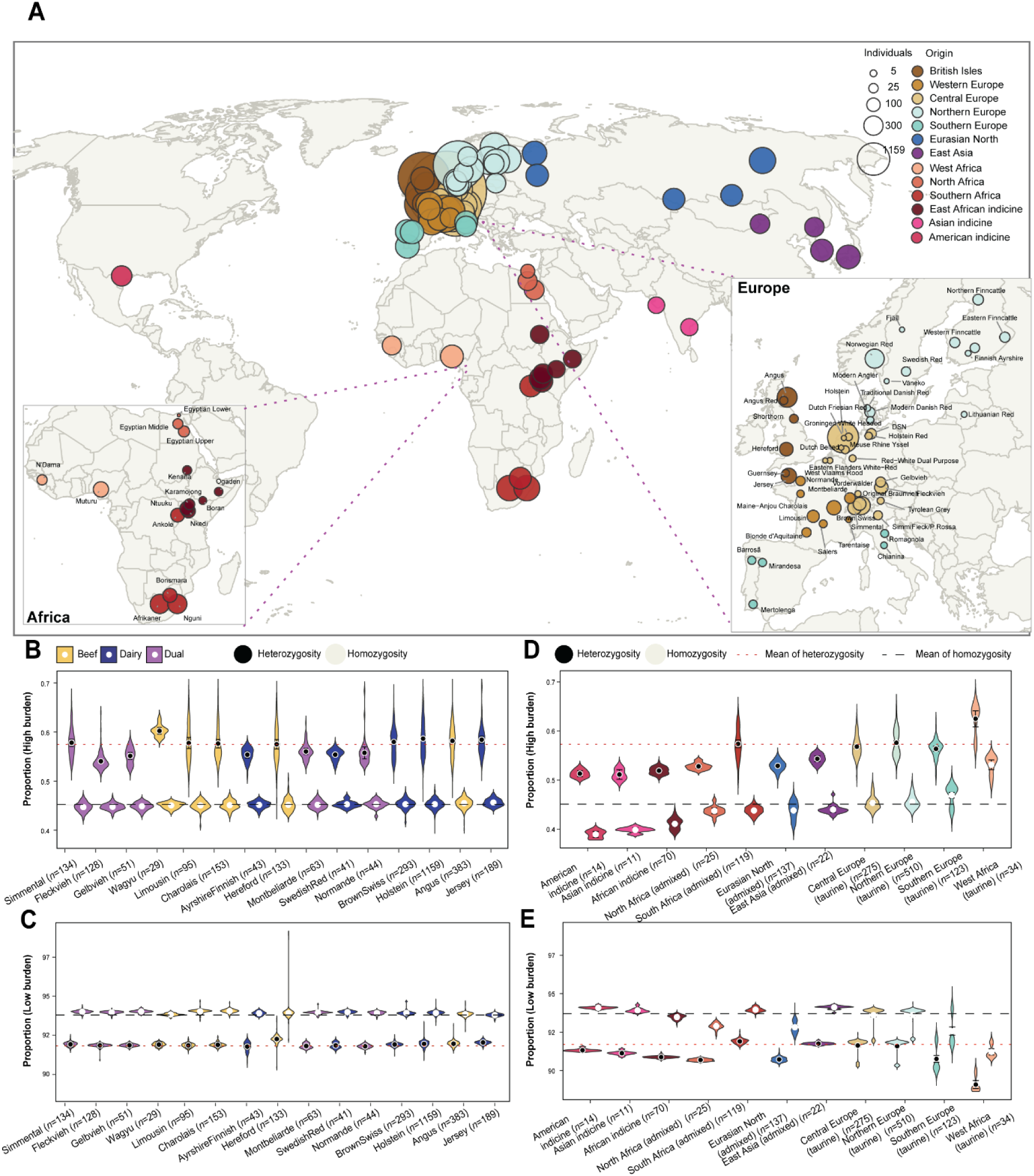
Population genetic load across cattle production systems and global regions. (A) Geographic origin of the cattle populations analysed. Point position denotes breed origin, size denotes the number of individuals sequenced, and color denotes geographic group. Insets show Europe and Africa in detail. All samples derive from the 1000 Bull Genomes Project and OPTIBOV. **(B)** Proportion of high-burden variants (BovCADD > 20) per individual (%) across commercial cattle breeds, grouped by production system (dairy, beef, and dual-purpose). Violin plots show the distributions of heterozygous and homozygous high-burden variants; points indicate the mean ± 1.96 standard errors. (C) As in (B), for low-burden variants (BovCADD < 5). (D) Proportion of high-burden variants (BovCADD > 20) per individual (%) across global cattle populations grouped by geographic region and subspecies. Points indicate the mean ± 1.96 standard errors. (E) As in (D), for low-burden variants (BovCADD < 5). Population definitions (**D,E**). All samples derive from the 1000 Bull Genomes Project and OPTIBOV. American indicine includes Brahman (14). Asian indicine includes Nelore (6) and Gir (5). East African indicine includes Nkedi (19), Ntuuku (19), Karamojong (11), Kenana (8), Ogaden (7), and Boran (6). North Africa includes Egyptian Upper (12), Egyptian Middle (10), and Egyptian Lower (3) cattle. Southern Africa includes Afrikaner (40), Bonsmara (20), Nguni (40) and Ankole (19). Eurasian North includes Yakut (44), Buryat (19), Altai (20), Kholmogory (32), and Yaroslavl (22). East Asia includes Menggu (11) and Yanbian (11). Central Europe includes Original Braunvieh (111), Deutsches Schwarzbuntes Niederungsrind (56), Meuse Rhine Yssel (25), Groningen White Headed (21), Tyrolean Grey (17), Tarentaise (12), Dutch Belted (11), Dutch Friesian Red (11), and West Vlaams Rood (11). Northern Europe includes Norwegian Red (345), Northern Finncattle (44), Swedish Red (41), Eastern Finncattle (40), and Western Finncattle (40). Southern Europe includes Mertolenga (29), Barrosã (28), Mirandesa (27), Romagnola (24), and Chianina (15). West Africa includes Muturu (24) and N’Dama (10). Global mean proportions (**A–D**) calculated across all individuals from the 1000 Bull Genomes Project and OPTIBOV, are shown as dashed reference lines (heterozygous high-burden = 0.57; homozygous high-burden = 0.45; heterozygous low-burden = 91.70; homozygous low-burden =93.70).

We next characterized load across geographic groups and subspecies (fig. S3, Table S7). West African taurine breeds carried the highest load in both classes and exceeded all commercial breeds (Muturu 0.542% homozygous and 0.642% heterozygous high-burden; N’Dama 0.505% and 0.583%), consistent with their reported low genetic diversity and historical bottlenecks (Fig. 5D, fig. S3A, S3B) [70]. European taurine populations were intermediate and exceeded crossbred populations, while indicine populations from Africa, the Americas, and Asia carried the lowest load (Brahman 0.389% homozygous high-burden; Gir 0.395%; Nelore 0.401%). The taurine–indicine contrast was the dominant axis of variation, with differences of 0.09–0.15 percentage points between subspecies compared with ∼0.01 within the European taurine group (Table S7). Within European taurine breeds, load tracked previously reported diversity patterns across local breeds (fig. S3, Table S7). Mirandesa, an Iberian local breed that recently underwent a bottleneck associated with intensive breeding [71], carried a significantly lower proportion of homozygous low-burden variants than the genetically diverse Barrosã and Mertolenga (91.74% versus 91.94% and 91.99%; FDR < 0.05), indicating reduced retention of benign variation (fig. S3C). Homozygous high-burden proportions were nominally higher in Mirandesa but did not differ significantly among the three breeds (fig. S3A, Table S6). Consistent with their elevated deleterious load, West African and European taurine populations also showed the lowest proportions of low-burden variants (Fig. 5E, fig. S3D, S3E, Table S6). Together, these patterns show that BovCADD-based genetic load captures population-specific differences in the accumulation and distribution of deleterious variation beyond broad effects of population size or subspecies background.

## DISCUSSION

Genetic variation in cattle is shaped by demographic history, admixture, selection, and genome architecture, which makes it difficult to distinguish deleterious variants from neutral background variation. Here we present BovCADD, a cattle-wide deleteriousness framework that improves variant interpretation and completion beyond limited single-source predictors and trait-specific prioritization tools. BovCADD integrates 88 functional annotations spanning evolutionary constraint, sequence context, regulatory and epigenetic activity, and gene and protein features, and provides precomputed scores for all 8.1 billion possible single-nucleotide substitutions in the bovine genome. It recapitulated functional gradients across genomic compartments and consequence classes, with high scores enriched among predicted loss-of-function and splice-disrupting variants while also resolving putatively functional variants in intronic and intergenic sequence, including at QTL-linked loci and their flanking regions. In validation, BovCADD separated pathogenic OMIA variants from background variation, and its top-ranked variants showed moderate concordance with trait-oriented prioritization frameworks such as FAETH [52] and Functional-And-Evolutionary Multi-trait Importance (FAEMI) [25]. Beyond SNVs prioritization, BovCADD supports analyses that require deleteriousness scores across the whole genome rather than at individual loci. Gene-based burden mapping identified genes accumulating rare deleterious variation and recovered coherent pathway-level enrichment, following approaches established for rare germline variation in human cohorts [53]. Applied within QTL intervals, the same framework distinguished putatively functional variants from background variation in both coding and regulatory sequence, extending trait-mapping resolution beyond the association signal itself. At the population level, stratifying variants by burden class allowed genetic load to be compared across breeds and geographic groups on a common scale.

To our knowledge, BovCADD is the first cattle-wide framework to combine variation from the largest available global whole-genome sequencing resources with a broad, multilayer annotation set for prioritizing deleterious SNVs. The model was trained and evaluated on 3,747 cattle genomes spanning 215 breeds from the 1000 Bull Genomes Project (Run 9) [72] and the LEAP-Agri OPTIBOV project [37, 73], encompassing *Bos taurus*, *Bos indicus*, and admixed populations across Europe, Africa, Asia, and the Americas. The feature set combines local sequence properties (e.g. GC/CpG, mutational class, DNA-shape [74], TSS proximity), multi-species conservation and constraint (phastCons, phyloP and GERP-derived scores [7]), and cattle regulatory activity derived from bovine Functional Annotation of Animal Genomes (FAANG) ATAC-seq/ChIP-seq maps and CTCF-defined chromatin elements [41], together with predicted TF motif distributions [45], consequence-based annotations (VEP [8], and SIFT [4]). Consistent with CADD-style frameworks in humans and other species, conservation-derived metrics provided the strongest single-annotation signal in every genomic region examined [9, 35]. This consistency has two sources. Conservation scores are defined at base-pair resolution genome-wide, whereas annotations such as GC content and the epigenetic marks are sparse or tissue-specific [75]. Second, conservation scores and the training labels measure closely related quantities: both are proxies for whether a site has been subject to selection [12]. However, sparse annotations contributed orthogonal information that conservation cannot capture, and their contributions varied by genomic context: coding-impact features such as SIFT dominated in missense sequence, whereas regulatory and sequence-context features contributed more in intronic, UTR, and intergenic regions. Reflecting this, 44.95% of substitutions scoring above BovCADD 20 were non-coding, indicating sensitivity beyond consequence class. Conversely, 65.59% of potential missense and 8.94% of potential stop-gained variants scored below 20, indicating that the model separates likely tolerated variants from more damaging ones within a single consequence class.

BovCADD also performed well in external validation, separating OMIA pathogenic variants from background variation with an AUC of 0.998, and ranking most catalogued OMIA variants among the highest-scoring substitutions genome-wide (score range 1.43–45.78; mean 22.32) [28]. Because a substantial fraction of high-scoring substitutions falls outside coding sequence, we further contrasted scores between coding and non-coding regions to establish a threshold for prioritizing deleterious non-coding variants. We then compared BovCADD with FAETH, a cattle-specific score optimized for trait-heritability prioritization across 34 commercially relevant traits [52]. The two frameworks differ in purpose, and their agreement reflected this: correlation was moderate among top-ranked variants but negligible across the remaining score range. This pattern is consistent with the way FAETH is constructed, since variants in the lower part of its distribution contribute little to the genomic relationship matrix from which the score is derived and are correspondingly less well resolved [9, 35, 52]. A further difference in scope is that FAETH is estimated from variants segregating in the sampled populations and is therefore restricted to the sites those samples contain, whereas BovCADD assigns a score to every possible substitution in the reference genome, including sites at which no variant has yet been observed. A related comparison comes from FAEMI, whose heat-stress ranking identified trait-associated variants whose human orthologs are predicted deleterious by human CADD [25]. Previous comparisons with human orthologs further support the need for BovCADD, as no cattle-specific deleteriousness score was available before this work [25]. Compared with these trait-specific resources, BovCADD does not require phenotype-specific training and can therefore be applied to loci and phenotypes that are absent from existing prioritization sets. By integrating signals from conservation, gene structure, coding impact and protein-level annotations, BovCADD adds resolution beyond trait-specific frameworks and provides a broadly applicable tool for variant interpretation.

Variant-level scores address one locus at a time, but most traits of economic and biological importance in cattle are polygenic, arising from many variants of small effect distributed across coding, transcription-factor binding, and regulatory regions of the genome [76, 77]. We therefore extended BovCADD from scoring individual substitutions to aggregating scores across loci, in three settings: the genome-wide distribution of rare deleterious variation, the burden carried by trait-relevant genes and QTL intervals, and population-level genetic load. Aggregating rare, high-scoring variants on a per-gene basis showed that deleterious burden is unevenly distributed across the genome in the 1000 Bull Genomes populations. This uneven distribution provides a starting point for more detailed analyses, including population-specific burden comparisons and pathway enrichment [53]. Two exemplar gene sets illustrate how gene-level burden tracks selection history. Among milk-protein genes, the breed carrying the lowest burden differed by locus: Holstein-Friesian at *CSN1S1*, *CSN2*, and *LALBA*, Jersey at *CSN1S2* and *CSN3*, which encodes the κ-casein governing milk coagulation and cheese-making properties [61]. At stress-response loci, burden separated temperate European taurine breeds from heat-adapted African and indicine populations, the latter carrying fewer high-burden variants [33]. These two patterns arise from different processes. Reduced burden in heat-adapted populations is consistent with sustained natural selection depleting deleterious variation. Artificial selection acts in the opposite direction genome-wide, and Holstein-Friesian and Jersey indeed carry the highest overall loads among commercial breeds, consistent with hitchhiking and reduced effective population size under intensive selection [78]. Yet at the loci under selection the direction reverses, with each breed carrying its lowest burden at the casein genes most relevant to the traits it has been bred for [54, 59, 60].

Genetic load reflects the expected reduction in fitness arising from the accumulation of deleterious mutations, either through recurrent mutation input or at mutation–selection balance [16, 79, 80]. Given the divergent subspecies backgrounds of cattle, we adopted a zygosity-stratified approach to genetic load analysis, building on traditional formulations of realized and masked load [16]. Partitioning load into homozygous and heterozygous components provides additional insight into the nature of deleterious variation carried by populations. Homozygous burden corresponds to the realized load, the fraction that is expressed and therefore exposed to purifying selection and is most informative about recessive deleterious alleles. Heterozygous burden corresponds to the masked load, which is shielded from selection in the heterozygous state and is expressed only as homozygosity increases [81]. By quantifying the proportions of high-burden and low-burden variants in both zygosity classes, BovCADD enables genetic load comparisons across breeds, subspecies, and geographic regions, and offers new insight into how genetic diversity, selective background, and demographic history interact. The elevated high-burden proportions in West African taurine breeds, and in the endangered Muturu in particular, are consistent with their reduced diversity, small effective population size, and documented bottlenecks [70, 82]. The lower burden in indicine populations conversely reflects their larger diversity and distinct selection history [83]. The elevated high burden in dairy breeds such as Jersey and Holstein supports the view that sustained intensive selection, combined with heavy use of elite sires, promotes accumulation of deleterious variation through hitchhiking and increased relatedness [84]. At a finer scale, the elevated homozygous load in Mirandesa is consistent with its recent bottleneck and intensive breeding history, while the lower loads in the more diverse Barrosã and Mertolenga follow the same logic [71]. The parallel depletion of low-burden variants in West African and commercial taurine populations reinforces the pattern and indicates that BovCADD-based burden metrics offer a workable framework for comparing the consequences of selection and demographic constraint across cattle populations.

Several limitations should be acknowledged. BovCADD uses L2-regularized logistic regression, a linear model that captures interactions only through explicitly constructed crossed features; we retained it for consistency with established CADD implementations and for interpretability [9–11], but did not test whether a nonlinear learner would extract more signal. The model also scores single-nucleotide substitutions only, excluding indels and structural variants that contribute substantially to phenotypic variation in cattle. Because the training contrast is defined by variant depletion relative to sequence-context-matched expectations, it is sensitive to regional variation in mutation rate, background selection, and GC-biased gene conversion [10], and the resulting score reflects evolutionary intolerance rather than trait effect size [9]. Finally, the scarcity of experimentally validated functional variants in cattle, particularly in non-coding sequence, limits benchmarking there. As in other species, CADD-style scores are best read as probabilistic indicators of deleteriousness that complement, rather than replace, association evidence and functional validation.

In conclusion, BovCADD is a genome-wide framework for identifying variants likely to be deleterious in cattle, trained without reference to any particular trait and applicable across coding and regulatory sequence alike. As genomic resources expand across commercial and indigenous breeds, the gap between variant discovery and functional interpretation will continue to widen. BovCADD narrows that gap by supplying a single ranked score for every possible substitution, reducing the need to generate, curate, and reconcile disparate functional datasets for each new question. We expect that publicly available precomputed scores will support rapid prioritization of candidate variants, functional annotation of QTL and GWAS intervals, genetic load assessment, and monitoring of the genetic basis of health, productivity, and adaptation in cattle populations worldwide.

## MATERIALS AND METHODS

### Sample collection

A total of 6,723 individuals representing 262 cattle breeds from the 1000 Bull Genomes Project (Run 9) [72], 533 individuals representing 25 breeds from the LEAP-Agri OPTIBOV project [37, 73] (https://subsites.wur.nl/en/optibov-project.html), and Muturu Genomic Resource [85] were combined to create a broad multi-breed cattle dataset. Because sequencing coverage in the OPTIBOV dataset was approximately 10×, and to ensure comparable data quality across datasets, a depth filter of >10 was applied to the 1000 Bull Genomes Project data [72], after which 3,747 individuals from 215 cattle breeds were retained for downstream analyses. Detailed information on sequencing coverage, sample identifiers, breed assignments, and release IDs is provided in Data S6. The final dataset included *Bos taurus*, *Bos indicus*, and admixed cattle populations from Europe, Africa, Asia, and the Americas, thereby capturing the major global cattle lineages and a wide range of demographic histories, breeding systems, and selection pressures. Sequencing data were generated at a mean depth of approximately 15× per individual, which provided sufficient resolution for robust variant discovery and genotype calling across diverse breeds. To maximize consistency across sources, Variant Call Format (VCF) files were merged and jointly processed using BCFtools v1.9 [86]. We then applied stringent filtering criteria, including read depth (DP) ≥ 4, variant quality (QUAL) ≥ 20, minor allele frequency (MAF) ≥ 0.01, and missing genotype rate ≤ 0.01, to retain high-confidence SNVs for subsequent model training and downstream analyses.

### Simulated and observed variants

Ancestral sequences were reconstructed from Ensembl EPO multi-species alignments encompassing 43 eutherian mammals (Ensembl release v113) [8]. The most recent common ancestor of cattle (*Bos taurus*; ARS-UCD1.2) and sheep (*Ovis aries*) were inferred from the alignment, representing the closest available ruminant outgroup. Sheep served as the outgroup to polarize each variant position as ancestral or derived relative to the bovine reference genome. Only positions with a confidently reconstructed ancestral state at the cattle-sheep ancestral node were retained for downstream analysis. This ancestral sequence served two distinct purposes: first, as the proxy-deleterious variants from the mutational background simulated data, independent of evolutionary constraint; and second, as the reference polarization framework for identifying observed bovine-derived substitutions (tolerated; proxy-benign) from 3,747 samples across 215 cattle breeds.

Simulated (proxy-deleterious) variant set: following the human and non-human CADD framework, we implemented a fully empirical simulator of sequence evolution adapted for the bovine genome. The simulator models single nucleotide substitutions incorporating trinucleotide context-dependent substitution rates, CpG-specific hypermutability, and local mutation-rate variation at approximately 1 Mb scales, consistent with the approach [9–11, 15]. Rate parameters and context effects were estimated directly from the reconstructed cattle-sheep ancestral sequence within the EPO 43-mammal alignment, capturing the substitution history along the bovine lineage. These parameters were then applied to the bovine reference genome (ARS-UCD1.2) to simulate genome-wide SNVs. Simulated variants were sampled to reflect the observed distribution of local sequence context and regional mutability, yielding a mutational background set equal in size to the observed variant set for balanced model training. To prevent contradictory labels in the training data, simulated variants were excluded at any position already represented in the observed set, defined as positions where a derived allele was present at DAF ≥ 95% in the combined 1000 Bull Genomes Project (Run 9) [72] and OPTIBOV project [37, 73].

Observed (proxy-benign) variant set: Bovine-derived substitutions were identified by contrasting the cattle reference genome (ARS-UCD1.2) with the reconstructed cattle-sheep ancestral sequence, enabling direct allele polarization at each callable site. Candidate derived sites were intersected with contemporary bovine population data from the 1000 Bull Genomes Project (Run 9) [72] and OPTIBOV project [37, 73]. Sites were retained where the derived allele frequency (DAF) was ≥ 95% across the combined cohort, capturing substitutions that are fixed or approaching fixation on the bovine lineage and therefore most likely to reflect variants that have persisted through the evolutionary filter. To ensure complete capture of near-fixed substitutions, we additionally included loci where the reference allele matched the inferred ancestral state, but the derived allele was observed at DAF ≥ 95% in the population data, representing cases where the reference genome retains the ancestral base despite near-complete substitution in the contemporary population. Applying these criteria yielded 41.9 million observed SNVs.

The final training set was assembled by pairing each observed SNV one-to-one with a context-matched simulated SNV, yielding a balanced dataset of 83.8 million variants in total. Observed variants served as the proxy-benign set, representing substitutions that have persisted and risen to high frequency under real evolutionary and selective pressures. Simulated variants served as the proxy-deleterious set, representing variants expected under the null model of sequence evolution in the absence of functional constraint. The contrast between these two sets provides the supervised learning signal from which the BovCADD model learns to distinguish tolerated variation from mutational background.

### Variant annotation matrix

Functional annotations for all single-nucleotide variants were generated using Ensembl VEP v113.3 with the ARS-UCD1.2 bovine reference genome and Ensembl gene models[8]. Core variant descriptors included chromosome, position, reference and alternate alleles, the transition–transversion indicator (isTv), and 13 VEP-derived consequence fields. For protein-coding variants, SIFT scores and categorical predictions were obtained via the VEP SIFT plugin to capture predicted amino acid–altering effects [4].

The 73 additional features capturing evolutionary, sequence-context, and regulatory information were incorporated from bovine-specific and comparative genomic resources (Table S2). Evolutionary conservation was quantified using phastCons and phyloP scores [6] derived from multiple sequence alignments spanning distinct phylogenetic depths, including mammalian alignments (UCSC multiz35way: cattle bosTau9, human hg38, mouse mm39, dog canFam4, pig susScr3, and opossum monDom5), Cetartiodactyla alignments (UCSC multiz35way: cattle bosTau9, sheep oviAri4, pig susScr3, and bottlenose dolphin turTru2), and ruminant-focused multi-species alignments (Ensembl EPO 43-mammal v113, including cattle *Bos taurus*, sheep *Ovis aries*, goat *Capra hircus*, zebu *Bos indicus*, and yak *Bos grunniens*), with the cattle sequence excluded from all conservation score calculations. Additional evolutionary constraint metrics were obtained from GERP single-site and element-level scores (gerpS, gerpN, gerpRS, gerpRSpval) computed from multi-mammal alignments [7].

Local sequence context features were derived directly from the reference genome and included GC and CpG content, transition–transversion class, DNA-shape parameters (HelT, MGW, ProT, Roll) [74], and TSS. Gene- and protein-level annotations included reference and derived amino acids VEP v113.3 [8], Grantham distance [46], SIFT score (SIFTcat and SIFTval) [4], and relative positions within the cDNA, CDS, and protein sequence VEP v113.3 [8].

Regulatory annotations were integrated from public bovine FAANG ATAC-seq and ChIP-seq datasets [41], representing open chromatin and histone modification landscapes across major tissues, including adipose, cerebellum, cortex, hypothalamus, liver, lung, muscle, spleen, neuronal cells and adult rumen (Table S1). Histone marks comprised H3K27ac, H3K27me3, H3K4me1, and H3K4me3, while CTCF ChIP-seq peaks were included to reflect chromatin insulation and loop boundary activity. Transcription Factor motif distribution scores were computed using binding profiles from the JASPAR CORE database [45] and scanned with FIMO (MEME Suite v5.5.9) [87]. Variants were annotated for overlap with these regulatory features’ peak regions to derive tissue-specific regulatory activity measures.

The resulting variant-by-annotation matrix comprised 88 features, capturing local sequence composition, evolutionary conservation, chromatin accessibility, regulatory activity, and protein-coding consequences. All figures were generated using ggplot2 v3.5.1 [88].

### Imputation

Some annotated features described above were excluded from model training because they were either uninformative for deleteriousness prediction or differed systematically between simulated variants and cattle–sheep ancestral substitutions due to technical annotation constraints (Table S2). These features were retained for variant identification or stratification purposes but were not used as predictors during model training. To ensure model robustness while preserving biological interpretability, missing values were handled using feature-specific imputation strategies based on data type, annotation source, and biological meaning. Metadata fields, including chromosome and genomic position, were excluded from imputation and used solely for variant identification. The isTv feature encodes whether a substitution represents a transversion (1) or a transition (0). Missing values, which arise primarily from ambiguous allele annotations, were imputed as 0.5, representing an uninformative midpoint between transition and transversion states. The isMasked feature indicates whether a variant falls within repeat-masked or low-complexity genomic regions. Missing values for this feature were imputed as 1, reflecting a conservative assumption that unannotated sites may reside in masked regions.

Categorical variables (e.g. reference and alternate alleles, variant consequence classes, amino acid states (oAA and nAA), protein domain annotations, and SIFT functional categories) were one-hot encoded, with missing values assigned to a dedicated “unknown” category (UD). This approach ensures that variants affected by incomplete annotation are retained in the training data while preventing the introduction of artificial signal or systematic bias toward either benign or deleterious classes.

Continuous sequence-context and evolutionary conservation features (e.g. GC content, CpG density, DNA shape parameters, and evolutionary constraint metrics (GERP scores, phyloP, and phastCons scores across mammalian, cetartiodactyl, and ruminant alignments)) were imputed using the genome-wide mean of each feature. Mean imputation reflects neutral baseline expectations and avoids inflating deleteriousness scores due to missing conservation annotations in poorly aligned or repetitive genomic regions. Distance-based features were imputed using biologically conservative constants. Distances to transcription start and end sites (minDistTSS and minDistTSE) were set to 100 kb when missing, representing distal regulatory contexts with minimal expected functional impact. Binary or count-based regulatory annotations derived from ATAC-seq, ChIP-seq, and TF motif overlap analyses were imputed with 0, corresponding to the absence of regulatory evidence. Predictors with sporadic missingness due to annotation gaps (e.g., Grantham scores, SIFT scores, and coding position metrics (positions within cDNA, CDS, and protein sequences)) were imputed with neutral default values (0). This choice preserves feature completeness while minimizing bias in deleteriousness inference arising from incomplete transcript or protein annotations.

### Model training

Variant deleteriousness was modeled using a supervised machine-learning framework based on logistic regression (LogisticRegression, scikit-learn), trained on bovine whole-genome variation. The training dataset comprised 41.9 million high-frequency bovine-derived alleles, representing variants retained across worldwide cattle populations, contrasted with 41.9 million simulated single-nucleotide variants, ensuring balanced class representation. Simulated variants were treated as the positive class (label = 1) and derived variants as the negative class (label = 0), following the CADD paradigm[9].

Each variant was represented by 88 genomic features integrating sequence context, evolutionary conservation, predicted protein impact, regulatory annotations, and positional information. Categorical annotations were encoded as Boolean indicator variables, and continuous features were standardized prior to model fitting. Model training was performed using five-fold cross-validation with an L2-regularized logistic regression classifier, using a regularization strength of C = 10.0 and a maximum of 100 iterations. In addition to the model trained on the full feature set, separate models were trained using predefined annotation subsets (Grantham, GERP-S, phyloP, phastCons, SIFT, and combined conservation features) to assess the contribution of individual feature classes at OMIA sites.

### Comparison of variant prioritization using FAETH

To compare BovCADD with an established cattle-specific functional annotation framework, we analyzed FAETH scores. FAETH quantifies the contribution of sequence variants with regulatory and evolutionary relevance to 34 commercially important bovine traits, including production, reproduction, management, and linear assessment traits [52]. The FAETH scores were originally generated on the UMD3.1.1 bovine reference genome (https://figshare.unimelb.edu.au/s/f42b718e81e63dc488ac). To enable direct comparison with BovCADD, which is based on ARS-UCD1.2, FAETH variant coordinates were converted to the ARS-UCD1.2 assembly using UCSC liftOver v2025 [89]. Of the 17,495,272 primary FAETH-scored variant sites, 17,136,973 were successfully mapped and retained for downstream analyses; for sites with sex-specific estimates, mean FAETH values across bulls and cows were used (Zenodo: https://zenodo.org/records/20667548).

FAETH scores were used to rank variants genome-wide for both the multibreed genomic relationship matrix (GRM) and within-breed GRMs. FAETH coordinates were lifted to ARS-UCD1.2 and retained only where reference and alternate alleles matched. High- and low-ranking FAETH variants were identified using the same rank-based strategy applied to BovCADD. Variants were assigned to rank-based bins (top 10%, 10–20%, through to the lowest 10%), and Pearson correlation between the two scores was calculated within each bin separately for each of the 29 autosomes. Correlations are reported as the median and interquartile range across chromosomes. Because BovCADD is trait-agnostic and FAETH is trait-informed, locus-focused comparisons were additionally performed within milk production–associated QTL regions obtained from the Animal QTLdb [30]. All figures were generated using ggplot2 v3.5.1 [88].

### Gene-based burden of rare germline variation

To quantify how deleterious variation accumulates across genes in cattle, we performed a genome-wide gene-based burden analysis using BovCADD on VCF data from the 1000 Bull Genomes Project (Run 9) [72], following principles established in large-scale human studies of rare germline variation [53]. Rare germline variants were filtered to retain rare alleles (MAF < 0.01, allele count > 3) and high-quality genotypes (DP > 4; mean sample coverage > 5) [53]. We restricted analyses to populations with ≥10 individuals and retained breeds with sufficient sample size, resulting in 4,570 individuals across 67 breeds (Data S6). Filtered variants were annotated by intersecting sites with precomputed BovCADD scores. Variant processing and filtering were performed using BCFtools v1.20[90] and intersected with precomputed BovCADD scores. Putatively deleterious variants were defined using multiple BovCADD thresholds (>20, >25, >30 and >35). Variants were then aggregated at the gene level based on the reference gene annotation. Genome-wide gene density patterns were visualized using circos plots generated with TBtools v2 [91]. Functional enrichment analysis of high-burden genes was performed using the Database for Annotation, Visualization, and Integrated Discovery (DAVID v2021) for Gene Ontology (GO) Biological Process and Kyoto Encyclopedia of Genes and Genomes (KEGG) terms, applying Benjamini–Hochberg multiple-testing correction (adjusted *P* < 0.05). All plots were generated in ggplot2 v3.5.1 [88].

### Gene-based burden of trait-relevant genes

To illustrate putative gene-based burden in functionally relevant regions, analyses focused on key genes and their ±2 kb flanking regions associated with milk-production and stress-response phenotypes, selected based on prior functional evidence [30, 32–34, 92]. Following established approaches, individual-level genetic load was calculated by weighting predicted variant effects according to zygosity [93]. For an individual *k*, genetic load was defined as:

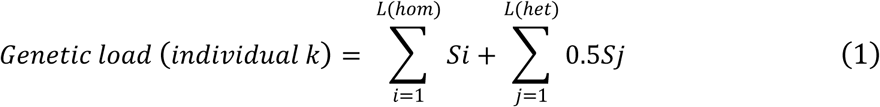

where *Si* and *Sj* denote the BovCADD score at homozygous and heterozygous loci, respectively, and *L(hom)* and *L(het)* represent the number of homozygous and heterozygous loci considered for individual *k*. When genetic load was compared across genomic regions of differing sizes, including QTLs and their flanking regions, scores were normalized by the total length of the analyzed region to obtain a length-normalized (weighted) genetic load. This normalization enables direct comparison of genetic loads across regions with different genomic extents. All figures were generated using ggplot2 v3.5.1 [88].

### Zygosity-stratified genetic load analysis

Genetic load refers to the reduction in fitness caused by the accumulation of deleterious mutations, either through recurrent mutation or at mutation–selection balance [16, 79]. Predicted variant-effect scores can be used to derive practical proxies of genetic load for individuals or populations by summing or averaging deleteriousness across the genome. Such metrics can be computed separately for homozygous and heterozygous genotypes and can be further restricted to variants that are fixed or segregated at high frequency within populations [16].

Because worldwide cattle populations have complex subspecies backgrounds and extensive admixture, direct comparisons of absolute counts of deleterious variants can be confounded by differences in overall variant discovery, heterozygosity, and demographic history. To improve comparability across breeds and lineages, we therefore quantified load as the proportion of sites falling into high-burden versus low-burden BovCADD classes within each zygosity state. Specifically, for each individual (or population), we calculated the fraction of homozygous and heterozygous genotyped sites that were high-burden (e.g., BovCADD > 20) and low-burden (e.g., BovCADD < 5), respectively, thereby standardizing for differences in the total number of genotypes available for analysis. Homozygous burden is particularly sensitive to recessive deleterious alleles and more directly reflects exposure of damaging variants, whereas heterozygous burden captures additive or partially recessive variants that may persist at low frequency. All figures were generated using ggplot2 v3.5.1 [88].

### Translating BovCADD across reference genome assemblies

BovCADD scores were computed on ARS-UCD1.2 (bosTau9; RefSeq GCF_002263795.1) and are reference-assembly dependent. Although ARS-UCD1.2 remains the most widely used bovine reference, newer studies increasingly use alternative assemblies such as ARS-UCD2.0 (RefSeq GCF_002263795.3). To enable the use of BovCADD across different bovine RefSeq, variant coordinates were lifted from ARS-UCD1.2 using UCSC liftOver v2025[89] with assembly-specific chain files (e.g., bosTau9ToGCF_002263795.3.over.chain.gz). For ARS-UCD1.3 (RefSeq GCF_002263795.2), we did not generate a separate resource and instead used ARS-UCD1.2 BovCADD directly because of ARS-UCD1.2-compatible coordinates. For ARS-UCD2.0, liftOver retained ∼99.97% of sites (∼7.9 billion SNVs), and the resulting per-assembly BovCADD tables are provided in Figshare (https://figshare.com/s/732b5e4306797bd66462).

## Acknowledgments

We thank the 1000 Bull Genomes Consortium for providing sequence data, and the breeders and breed associations who contributed samples.

## Author contributions

Conceptualization: J.G., R.C.

Methodology: J.G., M.D., J.S.

Formal analysis: J.G., M.D., J.S.

Investigation: Y.L., E.B.

Resources: C.G., J.K., N.G., D.K., M.M., R.C.

Data curation: J.G., M.D., J.S.

Writing – original draft: J.G.

Writing – review & editing: J.G., M.D., J.S., Y.L., E.B., C.G., J.K., N.G., D.K., M.M., M.G., H.B., R.C.

Supervision: M.D., M.G., H.B., R.C.

## Data availability

OPTIBOV whole-genome sequencing data for the indigenous European and African cattle analyzed in this study have been deposited in the European Nucleotide Archive (ENA) under accession numbers PRJEB90816 [94] (European) and PRJEB90914 [95] (African). Publicly available variant data were obtained from the 1000 Bull Genomes Project (Run 9) under PRJNA391427 [96] (ENA study ERZ14211345). Muturu data were obtained from the African Genomic Resource under PRJEB74565 [85]. Precomputed BovCADD scores for the ARS-UCD1.2 and ARS-UCD2.0 reference assemblies are publicly available on figshare (https://figshare.com/s/732b5e4306797bd66462). FAETH scores mapped to the ARS-UCD1.2 reference assembly were obtained from Zenodo (https://zenodo.org/records/20667548).

## Code availability

Custom code and workflows (Linux shell, R and Python) used for variant processing, BovCADD model training and scoring, and figure generation are available at https://github.com/junxingao888/BovCADD.

## Supplementary Materials

The PDF file includes:

Tables S1–S6.

## Other Supplementary Material for this manuscript includes the following

Data S1. Predictive performance of individual annotations. Receiver operating characteristic (ROC) area under the curve (AUC) values for each genomic annotation used in BovCADD, calculated by evaluating the ability of individual features to discriminate between derived and simulated variants.

Data S2. Correlation structure among genomic annotations. Pairwise correlation coefficients among the BovCADD annotations.

Data S3. Feature interaction weights in the regularized logistic regression model.

Data S4. Correlation between BovCADD and FAETH scores across score percentile bins.

Data S5. List of high-burden genes identified at thresholds of >20, >25, >30, and >35 rare deleterious variants per gene.

Data S6. Sample list for the 1000 Bull Genomes Project, BovCADD training data, OPTIBOV project, and samples used in gene-based burden analyses.

## REFERENCES

1. Porto-Neto LR, Kijas JW, Reverter A: The extent of linkage disequilibrium in beef cattle breeds using high-density SNP genotypes. Genet Sel Evol 2014, 46:22.

2. Goddard ME, Hayes BJ: Mapping genes for complex traits in domestic animals and their use in breeding programmes. Nat Rev Genet 2009, 10:381–391.

3. Andersson L, Archibald AL, Bottema CD, Brauning R, Burgess SC, Burt DW, Casas E, Cheng HH, Clarke L, Couldrey C, et al: Coordinated international action to accelerate genome-to-phenome with FAANG, the Functional Annotation of Animal Genomes project. Genome Biol 2015, 16:57.

4. Vaser R, Adusumalli S, Leng SN, Sikic M, Ng PC: SIFT missense predictions for genomes. Nat Protoc 2016, 11:1–9.

5. Xiang R, Berg Ivd, MacLeod IM, Hayes BJ, Prowse-Wilkins CP, Wang M, Bolormaa S, Liu Z, Rochfort SJ, Reich CM: Quantifying the contribution of sequence variants with regulatory and evolutionary significance to 34 bovine complex traits. Proceedings of the National Academy of Sciences 2019, 116:19398–19408.

6. Pollard KS, Hubisz MJ, Rosenbloom KR, Siepel A: Detection of nonneutral substitution rates on mammalian phylogenies. Genome Res 2010, 20:110–121.

7. Huber CD, Kim BY, Lohmueller KE: Population genetic models of GERP scores suggest pervasive turnover of constrained sites across mammalian evolution. PLoS Genet 2020, 16:e1008827.

8. McLaren W, Gil L, Hunt SE, Riat HS, Ritchie GR, Thormann A, Flicek P, Cunningham F: The ensembl variant effect predictor. Genome biology 2016, 17:1–14.

9. Kircher M, Witten DM, Jain P, O’Roak BJ, Cooper GM, Shendure J: A general framework for estimating the relative pathogenicity of human genetic variants. Nat Genet 2014, 46:310–315.

10. Rentzsch P, Witten D, Cooper GM, Shendure J, Kircher M: CADD: predicting the deleteriousness of variants throughout the human genome. Nucleic Acids Res 2019, 47:D886–d894.

11. Schubach M, Maass T, Nazaretyan L, Röner S, Kircher M: CADD v1.7: using protein language models, regulatory CNNs and other nucleotide-level scores to improve genome-wide variant predictions. Nucleic Acids Res 2024, 52:D1143–d1154.

12. Groß C, de Ridder D, Reinders M: Predicting variant deleteriousness in non-human species: applying the CADD approach in mouse. BMC Bioinformatics 2018, 19:373.

13. Fu C, van Schipstal J, Calus MPL, Duenk P: Genomic prediction using mCADD scores as prior information in a mouse population. Genetics 2025.

14. Groß C, Derks M, Megens HJ, Bosse M, Groenen MAM, Reinders M, de Ridder D: pCADD: SNV prioritisation in Sus scrofa. Genet Sel Evol 2020, 52:4.

15. Lensing K, van Schipstal JGC, de Ridder D, Groenen MAM, Derks MFL: A generic pipeline for CADD Score generation: chickenCADD and turkeyCADD. G3 (Bethesda) 2025.

16. Bertorelle G, Raffini F, Bosse M, Bortoluzzi C, Iannucci A, Trucchi E, Morales HE, van Oosterhout C: Genetic load: genomic estimates and applications in non-model animals. Nat Rev Genet 2022, 23:492–503.

17. Kim K, Kwon T, Dessie T, Yoo D, Mwai OA, Jang J, Sung S, Lee S, Salim B, Jung J: The mosaic genome of indigenous African cattle as a unique genetic resource for African pastoralism. Nature Genetics 2020, 52:1099–1110.

18. Felius M: Cattle breeds of the World. Brill; 2024.

19. Malek Ž, Romanchuk Z, Yaschun O, Jones G, Petersen J-E, Fritz S, See L: Improving the representation of cattle grazing patterns in the European Union. Environmental Research Letters 2024, 19:114077.

20. Godde CM, Mason-D’Croz D, Mayberry DE, Thornton PK, Herrero M: Impacts of climate change on the livestock food supply chain; a review of the evidence. Global food security 2021, 28:100488.

21. Derks MFL, Gross C, Lopes MS, Reinders MJT, Bosse M, Gjuvsland AB, de Ridder D, Megens HJ, Groenen MAM: Accelerated discovery of functional genomic variation in pigs. Genomics 2021, 113:2229–2239.

22. Fang L, Liu S, Liu M, Kang X, Lin S, Li B, Connor EE, Baldwin RLt, Tenesa A, Ma L, et al: Functional annotation of the cattle genome through systematic discovery and characterization of chromatin states and butyrate-induced variations. BMC Biol 2019, 17:68.

23. Li J, Kong N, Han B, Sul JH: Rare variants regulate expression of nearby individual genes in multiple tissues. PLoS Genet 2021, 17:e1009596.

24. Schulz H, Ruppert AK, Herms S, Wolf C, Mirza-Schreiber N, Stegle O, Czamara D, Forstner AJ, Sivalingam S, Schoch S, et al: Genome-wide mapping of genetic determinants influencing DNA methylation and gene expression in human hippocampus. Nat Commun 2017, 8:1511.

25. Xiang R, Breen E, Bolormaa S, Liu Z, Vander Jagt CJ, Dong M, Lindblad-Toh K, Rochfort S, Pryce JE, Chamberlain AJ, Goddard ME: Integrating extensive functional annotations and multiomics of cattle enhances climate resilience prediction and mapping. Proc Natl Acad Sci U S A 2025, 122:e2514736122.

26. Cheng JY, Stern AJ, Racimo F, Nielsen R: Detecting Selection in Multiple Populations by Modeling Ancestral Admixture Components. Mol Biol Evol 2022, 39.

27. Ongaro L, Mondal M, Flores R, Marnetto D, Molinaro L, Alarcón-Riquelme ME, Moreno-Estrada A, Mabunda N, Ventura M, Tambets K, et al: Continental-scale genomic analysis suggests shared post-admixture adaptation in the Americas. Hum Mol Genet 2021, 30:2123–2134.

28. Tammen I, Mather M, Leeb T, Nicholas FW: Online Mendelian Inheritance in Animals (OMIA): a genetic resource for vertebrate animals. Mamm Genome 2024, 35:556–564.

29. Cole JB: A simple strategy for managing many recessive disorders in a dairy cattle breeding program. Genet Sel Evol 2015, 47:94.

30. Hu ZL, Park CA, Reecy JM: Bringing the Animal QTLdb and CorrDB into the future: meeting new challenges and providing updated services. Nucleic Acids Res 2022, 50:D956–d961.

31. Comin A, Cassandro M, Chessa S, Ojala M, Dal Zotto R, De Marchi M, Carnier P, Gallo L, Pagnacco G, Bittante G: Effects of composite β-and κ-casein genotypes on milk coagulation, quality, and yield traits in Italian Holstein cows. Journal of dairy science 2008, 91:4022–4027.

32. Ma Y, Khan MZ, Xiao J, Alugongo GM, Chen X, Chen T, Liu S, He Z, Wang J, Shah MK: Genetic markers associated with milk production traits in dairy cattle. Agriculture 2021, 11:1018.

33. Rakib M, Messina V, Gargiulo J, Lyons N, Garcia S: Graduate student literature review: Potential use of hsp70 as an indicator of heat stress in dairy cows—A review. Journal of Dairy Science 2024, 107:11597–11610.

34. Zeng L, Qu K, Zhang J, Huang B, Lei C: Genes related to heat tolerance in cattle—A review. Animal Biotechnology 2023, 34:1840–1848.

35. Groß C, Derks M, Megens H-J, Bosse M, Groenen MA, Reinders M, De Ridder D: pCADD: SNV prioritisation in Sus scrofa. Genetics Selection Evolution 2020, 52:4.

36. Hayes BJ, Daetwyler HD: 1000 Bull Genomes Project to Map Simple and Complex Genetic Traits in Cattle: Applications and Outcomes. Annu Rev Anim Biosci 2019, 7:89–102.

37. Ginja C, Gao J, Kantanen J, Ghanem N, Kugonza D, Makgahlela M, Pires AE, Usié A, Dibbits B, de Sousa CB, et al: Whole genome sequences of 289 native cattle from Finland, the Netherlands, and Portugal. Sci Data 2025, 12:1897.

38. McLaren W, Gil L, Hunt SE, Riat HS, Ritchie GR, Thormann A, Flicek P, Cunningham F: The ensembl variant effect predictor. Genome biology 2016, 17:122.

39. Kuhn RM, Haussler D, Kent WJ: The UCSC genome browser and associated tools. Briefings in bioinformatics 2013, 14:144–161.

40. Wang J, Yuan W, Liu F, Liu G, Geng X, Li C, Zhang C, Li N, Li X: Epigenetic basis for the establishment of ruminal tissue-specific functions in bovine fetuses and adults. J Genet Genomics 2025, 52:78–92.

41. Kern C, Wang Y, Xu X, Pan Z, Halstead M, Chanthavixay G, Saelao P, Waters S, Xiang R, Chamberlain A, et al: Functional annotations of three domestic animal genomes provide vital resources for comparative and agricultural research. Nat Commun 2021, 12:1821.

42. de Souza MM, Zerlotini A, Geistlinger L, Tizioto PC, Taylor JF, Rocha MIP, Diniz WJS, Coutinho LL, Regitano LCA: A comprehensive manually-curated compendium of bovine transcription factors. Sci Rep 2018, 8:13747.

43. Cooper GM, Stone EA, Asimenos G, Green ED, Batzoglou S, Sidow A: Distribution and intensity of constraint in mammalian genomic sequence. Genome Res 2005, 15:901–913.

44. Siepel A, Bejerano G, Pedersen JS, Hinrichs AS, Hou M, Rosenbloom K, Clawson H, Spieth J, Hillier LW, Richards S, et al: Evolutionarily conserved elements in vertebrate, insect, worm, and yeast genomes. Genome Res 2005, 15:1034–1050.

45. Rauluseviciute I, Riudavets-Puig R, Blanc-Mathieu R, Castro-Mondragon JA, Ferenc K, Kumar V, Lemma RB, Lucas J, Chèneby J, Baranasic D, et al: JASPAR 2024: 20th anniversary of the open-access database of transcription factor binding profiles. Nucleic Acids Res 2024, 52:D174–d182.

46. Grantham R: Amino acid difference formula to help explain protein evolution. Science 1974, 185:862–864.

47. Ng PC, Henikoff S: SIFT: Predicting amino acid changes that affect protein function. Nucleic Acids Res 2003, 31:3812–3814.

48. Zou F, Wang Y, Yang Y, Zhou K, Chen Y, Song J: Supervised feature learning via L2-norm regularized logistic regression for 3D object recognition. Neurocomputing 2015, 151:603–611.

49. Ewing B, Green P: Base-calling of automated sequencer traces using phred. II. Error probabilities. Genome Res 1998, 8:186–194.

50. Liu Y, Bijl E, Gao J, Gonzalez-Prendes R, Groenen MA, Kantanen J, Ginja C, Ghanem N, Kugonza DR, Makgahlela M: Global analysis of bovine milk protein variants using multi-breed DNA sequence data. BMC Genomics 2026.

51. Teng J, Wang D, Zhao C, Zhang X, Chen Z, Liu J, Sun D, Tang H, Wang W, Li J, et al: Longitudinal genome-wide association studies of milk production traits in Holstein cattle using whole-genome sequence data imputed from medium-density chip data. J Dairy Sci 2023, 106:2535–2550.

52. Xiang R, Berg IVD, MacLeod IM, Hayes BJ, Prowse-Wilkins CP, Wang M, Bolormaa S, Liu Z, Rochfort SJ, Reich CM, et al: Quantifying the contribution of sequence variants with regulatory and evolutionary significance to 34 bovine complex traits. Proc Natl Acad Sci U S A 2019, 116:19398–19408.

53. Ivarsdottir EV, Gudmundsson J, Tragante V, Sveinbjornsson G, Kristmundsdottir S, Stacey SN, Halldorsson GH, Magnusson MI, Oddsson A, Walters GB, et al: Gene-based burden tests of rare germline variants identify six cancer susceptibility genes. Nat Genet 2024, 56:2422–2433.

54. Caroli AM, Chessa S, Erhardt GJ: Invited review: milk protein polymorphisms in cattle: effect on animal breeding and human nutrition. J Dairy Sci 2009, 92:5335–5352.

55. Singh MK, Shin Y, Han S, Ha J, Tiwari PK, Kim SS, Kang I: Molecular Chaperonin HSP60: Current Understanding and Future Prospects. Int J Mol Sci 2024, 25.

56. Lee-Yoon D, Easton D, Murawski M, Burd R, Subjeck JR: Identification of a major subfamily of large hsp70-like proteins through the cloning of the mammalian 110-kDa heat shock protein. Journal of Biological Chemistry 1995, 270:15725–15733.

57. Schuermann JP, Jiang J, Cuellar J, Llorca O, Wang L, Gimenez LE, Jin S, Taylor AB, Demeler B, Morano KA, et al: Structure of the Hsp110:Hsc70 nucleotide exchange machine. Mol Cell 2008, 31:232–243.

58. Rong Y, Zeng M, Guan X, Qu K, Liu J, Zhang J, Chen H, Huang B, Lei C: Association of HSF1 Genetic Variation with Heat Tolerance in Chinese Cattle. Animals (Basel) 2019, 9.

59. Prinzenberg EM, Weimann C, Brandt H, Bennewitz J, Kalm E, Schwerin M, Erhardt G: Polymorphism of the bovine CSN1S1 promoter: linkage mapping, intragenic haplotypes, and effects on milk production traits. J Dairy Sci 2003, 86:2696–2705.

60. Brew K, Vanaman TC, Hill RL: The role of alpha-lactalbumin and the A protein in lactose synthetase: a unique mechanism for the control of a biological reaction. Proc Natl Acad Sci U S A 1968, 59:491–497.

61. Bonfatti V, Giantin M, Gervaso M, Coletta A, Dacasto M, Carnier P: Effect of CSN1S1-CSN3 (α(S1)-κ-casein) composite genotype on milk production traits and milk coagulation properties in Mediterranean water buffalo. J Dairy Sci 2012, 95:3435–3443.

62. Gambra R, Peñagaricano F, Kropp J, Khateeb K, Weigel KA, Lucey J, Khatib H: Genomic architecture of bovine κ-casein and β-lactoglobulin. J Dairy Sci 2013, 96:5333–5343.

63. Gao J, Ginja C, Liu Y, Kantanen J, Ghanem N, Kugonza D, Makgahlela M, Okwasiimire R, Bovenhuis H, Groenen MAM, Crooijmans R: Distinct adaptation and ancestral retention signals in African and European indigenous cattle genomes. Commun Biol 2026, 9.

64. Gifford-Gonzalez D, Hanotte O: Domesticating animals in Africa: implications of genetic and archaeological findings. Journal of World Prehistory 2011, 24:1–23.

65. Fuller DQ, Boivin N: Crops, cattle and commensals across the Indian Ocean. Current and potential archaeobiological evidence. Études Océan Indien 2009:13–46.

66. Bollongino R, Burger J, Powell A, Mashkour M, Vigne JD, Thomas MG: Modern taurine cattle descended from small number of near-eastern founders. Mol Biol Evol 2012, 29:2101–2104.

67. Chen S, Lin BZ, Baig M, Mitra B, Lopes RJ, Santos AM, Magee DA, Azevedo M, Tarroso P, Sasazaki S, et al: Zebu cattle are an exclusive legacy of the South Asia neolithic. Mol Biol Evol 2010, 27:1–6.

68. Loftus RT, MacHugh DE, Bradley DG, Sharp PM, Cunningham P: Evidence for two independent domestications of cattle. Proceedings of the National Academy of Sciences 1994, 91:2757–2761.

69. Salehi R, Javanmard A, Mokhber M, Alijani S: Estimating Linkage Disequilibrium and Effective Population Size Across Generations in Holstein Cattle. Vet Med Sci 2025, 11:e70684.

70. Mauki DH, Tijjani A, Ma C, Ng’ang’a SI, Mark AI, Sanke OJ, Abdussamad AM, Olaogun SC, Ibrahim J, Dawuda PM: Genome-wide investigations reveal the population structure and selection signatures of Nigerian cattle adaptation in the sub-Saharan tropics. BMC genomics 2022, 23:306.

71. da Fonseca RR, Ureña I, Afonso S, Pires AE, Jørsboe E, Chikhi L, Ginja C: Consequences of breed formation on patterns of genomic diversity and differentiation: the case of highly diverse peripheral Iberian cattle. BMC Genomics 2019, 20:334.

72. Hayes BJ, Daetwyler HD: 1000 bull genomes project to map simple and complex genetic traits in cattle: applications and outcomes. Annual review of animal biosciences 2019, 7:89–102.

73. Gao J, Gonzalez-Prendes R, Liu Y, Kantanen J, Ginja C, Ghanem N, Kugonza DR, Makgahlela M, Bovenhuis H, Groenen MAM, Crooijmans R: Evidence of early genomic selection in Holstein Friesian across African and European ecosystems. BMC Genomics 2025, 26:615.

74. Li J, Rohs R: Deep DNAshape webserver: prediction and real-time visualization of DNA shape considering extended k-mers. Nucleic Acids Res 2024, 52:W7–w12.

75. Lensing K, van Schipstal JGC, de Ridder D, Groenen MAM, Derks MFL: A generic pipeline for CADD score generation: chickenCADD and turkeyCADD. G3 Genes|Genomes|Genetics 2026, 16.

76. Crouch DJM, Bodmer WF: Polygenic inheritance, GWAS, polygenic risk scores, and the search for functional variants. Proc Natl Acad Sci U S A 2020, 117:18924–18933.

77. Niu Q, Zhang T, Xu L, Wang T, Wang Z, Zhu B, Zhang L, Gao H, Song J, Li J, Xu L: Integration of selection signatures and multi-trait GWAS reveals polygenic genetic architecture of carcass traits in beef cattle. Genomics 2021, 113:3325–3336.

78. Makino T, Rubin CJ, Carneiro M, Axelsson E, Andersson L, Webster MT: Elevated Proportions of Deleterious Genetic Variation in Domestic Animals and Plants. Genome Biol Evol 2018, 10:276–290.

79. Agrawal AF, Whitlock MC: Mutation load: the fitness of individuals in populations where deleterious alleles are abundant. Annual Review of Ecology, Evolution, and Systematics 2012, 43:115–135.

80. Henn BM, Botigué LR, Bustamante CD, Clark AG, Gravel S: Estimating the mutation load in human genomes. Nature Reviews Genetics 2015, 16:333–343.

81. Dussex N, Morales HE, Grossen C, Dalén L, van Oosterhout C: Purging and accumulation of genetic load in conservation. Trends in ecology & evolution 2023, 38:961–969.

82. Tijjani A, Utsunomiya YT, Ezekwe AG, Nashiru O, Hanotte O: Genome Sequence Analysis Reveals Selection Signatures in Endangered Trypanotolerant West African Muturu Cattle. Front Genet 2019, 10:442.

83. Mavunga TK, Sölkner J, Mészáros G, Pichler R, Zorobouragui L, Traore A, Tapsoba ASR, Hamed H, Hassan YA, Chileshe B, et al: Genome-wide analysis reveals differential admixture dynamics and historical demographic contractions in African cattle. Sci Rep 2026, 16:6495.

84. Brito LF, Bedere N, Douhard F, Oliveira HR, Arnal M, Peñagaricano F, Schinckel AP, Baes CF, Miglior F: Review: Genetic selection of high-yielding dairy cattle toward sustainable farming systems in a rapidly changing world. Animal 2021, 15 Suppl 1:100292.

85. ENA European Nucleotide Archive. [https://identifiers.org/ena.embl:PRJEB74565]

86. Li H: A statistical framework for SNP calling, mutation discovery, association mapping and population genetical parameter estimation from sequencing data. Bioinformatics 2011, 27:2987–2993.

87. Bailey TL, Boden M, Buske FA, Frith M, Grant CE, Clementi L, Ren J, Li WW, Noble WS: MEME SUITE: tools for motif discovery and searching. Nucleic Acids Res 2009, 37:W202–208.

88. Wickham H: ggplot2: elegant graphics for data analysis Springer-Verlag New York; 2009. Preprint at 2016, 2:15545–15550.

89. Perez G, Barber GP, Benet-Pages A, Casper J, Clawson H, Diekhans M, Fischer C, Gonzalez JN, Hinrichs AS, Lee CM, et al: The UCSC Genome Browser database: 2025 update. Nucleic Acids Res 2025, 53:D1243–d1249.

90. Danecek P, Bonfield JK, Liddle J, Marshall J, Ohan V, Pollard MO, Whitwham A, Keane T, McCarthy SA, Davies RM: Twelve years of SAMtools and BCFtools. Gigascience 2021, 10:giab008.

91. Chen C, Chen H, Zhang Y, Thomas HR, Frank MH, He Y, Xia R: TBtools: an integrative toolkit developed for interactive analyses of big biological data. Molecular plant 2020, 13:1194–1202.

92. Huang W, Peñagaricano F, Ahmad KR, Lucey JA, Weigel KA, Khatib H: Association between milk protein gene variants and protein composition traits in dairy cattle. J Dairy Sci 2012, 95:440–449.

93. Speak SA, Birley T, Bortoluzzi C, Clark MD, Percival-Alwyn L, Morales HE, van Oosterhout C: Genomics-informed captive breeding can reduce inbreeding depression and the genetic load in zoo populations. Mol Ecol Resour 2024, 24:e13967.

94. ENA European Nucleotide Archive [https://identifiers.org/ena.embl:PRJEB90816]

95. ENA European Nucleotide Archive. [https://identifiers.org/ena.embl:PRJEB90914]

96. ENA European Nucleotide Archive. [https://identifiers.org/ena.embl:PRJNA391427]

